# The chromosome-scale genome of the pine bark adelgid reveals conserved macrosynteny across Adelgidae

**DOI:** 10.64898/2026.08.10.743978

**Authors:** Dustin T. Dial, Kaitlyn N. Camp, Bryan M.T. Brunet, Carol D. von Dohlen, Gaelen R. Burke, Nathan P. Havill

## Abstract

Chromosome evolution varies widely across Aphidomorpha: several aphid lineages exhibit extensive interautosomal reshuffling, whereas comparisons involving grape phylloxera and *Adelges* suggest greater chromosome conservation outside Aphididae. However, all published adelgid genomes represent the genus *Adelges*, leaving conservation across deeper adelgid divergences unresolved. Here, we report a chromosome-scale genome for the pine bark adelgid, *Pineus strobi*, providing the first genome for the genus *Pineus* and extending genomic sampling to an early-diverging adelgid lineage. Synteny analyses revealed broad conservation of major linkage groups between *Pineus* and *Adelges* across approximately 90 million years. The five grape phylloxera chromosomes also showed broad correspondence to the ten adelgid chromosomes, consistent with a limited number of chromosomal fusions or fissions and relatively little exchange among major linkage groups. These findings strengthen evidence that such extensive interautosomal reshuffling is not characteristic of Aphidomorpha as a whole. We also recovered a complete 25.3 kb mitochondrial genome with expanded, repeat-rich noncoding regions and complete circular genomes for the obligate nutritional symbionts “*Candidatus* Annandia pinicola” and “*Candidatus* Hartigia pinicola.” Comparisons with *Pineus similis* symbionts revealed conserved coding capacity and genome-wide synteny, supporting conservation of this dual nutritional symbiosis in pine-associated adelgids. Finally, we recovered the first complete *Wolbachia* genome reported from an adelgid. Because *Wolbachia* can induce parthenogenesis, its presence in *P. strobi* raises the possibility that it contributes to the maintenance or reinforcement of parthenogenesis in a species lacking a viable sexual generation. Together, these genomes provide an integrated resource for aphidomorph evolution and symbiosis.

**Significance statement:** Several aphid lineages exhibit extensive reshuffling among autosomes, whereas previous comparisons with grape phylloxera and *Adelges cooleyi* suggested greater chromosome conservation outside Aphididae; however, all published adelgid genomes have represented *Adelges*, leaving the long-term stability of chromosome organization across Adelgidae unresolved. The first chromosome-scale genome from *Pineus* reveals broad conservation of major linkage groups across approximately 90 million years of adelgid evolution and clear correspondence with the five chromosomes of the grape phylloxera, supporting chromosome conservation outside Aphididae. The accompanying mitochondrial and symbiont genomes provide an integrated resource for studying aphidomorph genome evolution and symbiosis.

## Introduction

Adelgids are conifer-feeding insects within Aphidomorpha, a clade that also includes aphids (Aphididae) and phylloxerans (Phylloxeridae). Adelgidae is a comparatively species-poor group with two genera, *Adelges* and *Pineus* (Favret 2026) and includes several invasive forest pests, including the hemlock woolly adelgid, *Adelges tsugae*, and balsam woolly adelgid, *Adelges piceae* (Havill and Foottit 2007). Adelgids exhibit complex life cycles, host-plant specialization, gall formation, and obligate nutritional symbioses, making them important systems for studying the evolution of host-plant and host-microbe interactions (Havill et al. 2007; Sano and Ozaki 2012; von Dohlen et al. 2013; von Dohlen et al. 2017; Weglarz et al. 2018; Mech et al. 2019; Dial et al. 2022). However, genomic resources for adelgids remain limited relative to aphids, restricting comparative analyses across Aphidomorpha.

Chromosome-scale genomes are particularly important for understanding genome evolution in Aphidomorpha. Aphids show extensive autosomal rearrangement and long-term conservation of X chromosome gene content, but how this contrasts with adelgids is still relatively unexplored (Mathers et al. 2021; Li et al. 2023; Huang et al. 2025). Recent chromosome-scale assemblies for *Adelges tsugae* and *Adelges abietis*, along with the high-quality draft genome of *Adelges cooleyi*, have begun to establish adelgids as a comparative genomic system (Dial et al. 2023; Brunet et al. 2025; Glendening et al. 2026). These studies have shown few genome rearrangements in adelgids, but available high-quality adelgid genomes remain concentrated in *Adelges*. Genomic resources from *Pineus*, the *Pinus*-associated lineage recovered as sister to the remaining sampled adelgid lineages (Havill et al. 2007; Dial et al. 2023), are lacking and would allow testing to determine whether patterns observed in *Adelges* reflect broader features of adelgid genome evolution.

The pine bark adelgid, *Pineus strobi*, is native to eastern North America, where it is primarily associated with eastern white pine, *Pinus strobus* (Fig. 1) (Raske and Hudson 1964; Wantuch et al. 2017; Darr et al. 2018). It typically reproduces parthenogenetically through sequential generations of wingless females feeding on the bark, branches, and bases of needle whorls. Although *P. strobi* can produce winged sexuparae that migrate to black spruce, *Picea mariana*, offspring on spruce have not been observed to complete development, resulting in an effectively anholocyclic life cycle on white pine (Raske and Hudson 1964; Darr et al. 2018). *Pineus strobi* is generally not considered a major pest, but it can reach high local densities and has been reported outside its native eastern North American range, including western North America (Darr et al. 2018) and Europe (Carter 1971). Like other adelgids, *P. strobi* hosts bacteriocyte-associated bacterial symbionts, including “*Candidatus* Annandia pinicola” and “*Candidatus* Hartigia pinicola,” making it relevant for comparative studies of adelgid nutritional symbioses and symbiont genome evolution (Toenshoff et al. 2014; Dial et al. 2022).

**Figure 1.**
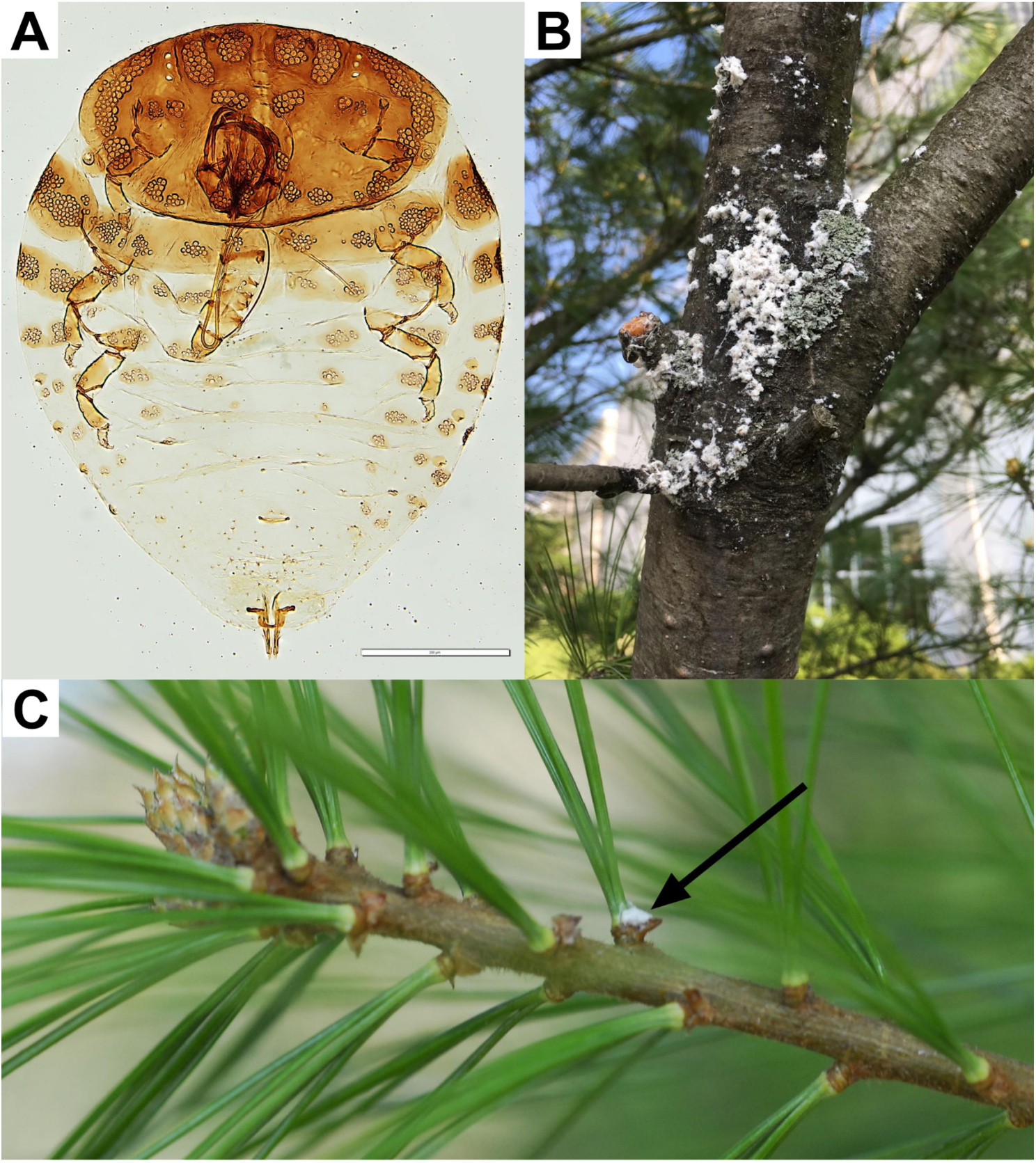
Morphology and host association of the pine bark adelgid, *Pineus strobi*. A) Cleared whole-mount specimen of an adult *P. strobi* collected in Foscoe, North Carolina, USA (Yale Peabody Museum accession ENT995701). B) Aggregations of wax-covered adelgids on pine bark in Ithaca, New York, USA. C) Wax-covered individual on a pine shoot (indicated by the black arrow) in Hamden, Connecticut, USA.

Here, we present a chromosome-scale genome assembly and associated endosymbiont and mitochondrial genomes for *P. strobi*. Comparative synteny analysis with available adelgid genomes supports broad conservation of macrosynteny among currently sequenced adelgids, contrasting with the extensive autosomal reshuffling observed in aphids. This genome provides the first chromosome-scale reference for *Pineus* and expands the phylogenetic breadth of genomic resources available for studying chromosome evolution, phylogeny, symbiosis, and diversification in Adelgidae.

## Results and Discussion

### Genome characteristics of *Pineus strobi*

We generated a chromosome-scale genome assembly for *P. strobi* using PacBio HiFi and Arima Hi-C sequencing. The final assembly spanned 217.47 Mb across 147 scaffolds, with a scaffold N50 of 18.63 Mb and a longest scaffold of 32.56 Mb (Table 1). Ten chromosome-scale scaffolds accounted for 94% of the assembly, consistent with a prior karyotype report of ten chromosomes in *P. strobi* (Steffan 1968; Blackman and Eastop 1994). The assembled genome size was similar to the k-mer-based estimate of 223.75 Mb (Supplementary Fig. S1) and a mean flow cytometric estimate of 281.8 ± 1.8 Mbp (n=3). Assembly completeness was high, with BUSCO recovering 99.1% complete Hemiptera odb10 orthologs, including 97.2% single-copy and 2.0% duplicated BUSCOs, with only 0.2% fragmented and 0.7% missing (Table 1). Merqury estimated a consensus quality value of QV 64.01, corresponding to an estimated base-error rate of 3.97 × 10⁻⁷. The PacBio HiFi assembly was already highly contiguous before Hi-C scaffolding, with a contig N50 of 16.73 Mb, 153 contigs in total, and a longest contig of 26.22 Mb. Hi-C data from an independently collected conspecific sample were then used to order and orient contigs into chromosome-scale pseudomolecules. These Hi-C reads showed high concordance with the PacBio HiFi assembly, with a samtools stats alignment identity of 99.27% across mapped bases. The Hi-C contact map showed strong, continuous diagonal interaction signal within each of the ten chromosome-scale scaffolds, supporting their final ordering and orientation (Supplementary Fig. S2).

**Table 1.** Assembly and annotation statistics of *P. strobi*.

| Characteristics | Values |
| --- | --- |
| **<u>Genome assembly</u>** |  |
| Assembly size (Mb) | 217.47 |
| Number of chromosomes | 10 |
| Number of scaffolds/contigs | 147/153 |
| Longest scaffold/contig (Mb) | 32.56/26.22 |
| N50 scaffold/contig length | 18.63/16.73 |
| GC content (%) | 31.3 |
| BUSCO completeness (%) | C:99.1%[S:97.2%,D:2.0%], F:0.2%, M:0.7% |
| **<u>Repetitive elements</u>** |  |
| DNA transposons | 2,988 (0.51%) |
| LINEs | 23,179 (3.75%) |
| LTRs | 808 (0.23%) |
| SINEs | 0 (0.00%) |
| Simple repeats | 121,189 (2.15%) |
| Unclassified | 94,260 (17.22%) |
| Total masked sequence | 53.04 Mb (24.39%) |
| **<u>Gene annotation</u>** |  |
| Total genes | 10,758 |
| Protein coding genes | 9,805 |
| Transcripts | 13,079 |
| Transcript variants | 2,179 |
| BUSCO completeness (%) | C: 99.0%[S:96.8%,D:2.2%], F:0.0%, M:1.0% |

EGAPx identified 10,758 genes, including 9,805 protein-coding genes, 672 noncoding genes, and 256 non-transcribed pseudogenes (Table 1). EGAPx predicted 13,079 mRNA transcripts, with transcript variants identified for 2,179 genes. BUSCO analysis of the EGAPx-predicted proteome recovered 99.0% of the 2,510 Hemiptera odb10 orthologs as complete, including 96.8% single-copy and 2.2% duplicated BUSCOs, with no fragmented BUSCOs and 1.0% missing, indicating a highly complete annotation. RepeatMasker annotated 53.04 Mb of repetitive sequence in the *P. strobi* assembly, corresponding to 24.39% of the genome (Table 1). The repeat landscape was dominated by unclassified repeats, which accounted for 37.46 Mb and represented approximately 70.6% of all repeat-masked sequences. Among classified transposable elements, LINEs were the largest component, whereas LTR elements, DNA transposons, SINEs, satellites, and rolling-circle elements were rare or undetected. Comparison with other sequenced adelgid and phylloxerid genomes showed that the *P. strobi* genome is relatively compact, with genome size and predicted coding content more similar to *A. tsugae* than to *A. cooleyi* or *A. abietis* (Table 2). Although the sample sizes are low, these differences are consistent with genome expansion along the lineage leading to larch- and Douglas-fir-associated adelgids, potentially involving increases in both repetitive sequence and annotated coding content.

**Table 2.** Comparison of genome statistics across adelgids and phylloxerids.

|  | Genome size | Chromosomes | CDS | Repeat percent |
| --- | --- | --- | --- | --- |
| *Pineus strobi* | 217.47 | 10 | 9,805 | 24.39 |
| *Adelges tsugae* | 216.56 | 10 | 11,800 | 17.90 |
| *Adelges cooleyi* | 270.2 | 11<sup>a</sup> | 13,556 | 43.67<sup>b</sup> |
| *Adelges abietis* | 253.16 | 10 | 12,060 | 28.31 |
| *Daktulosphaira vitifoliae* | 282.86 | 5 | 15,582 | 53.26<sup>b</sup> |

<sup>a</sup>Scaffolds have not been assigned to chromosomes
<sup>b</sup>Repeats were masked with WindowMasker. All others were masked with RepeatMasker and Repeatmodeler.

Terminal repeat analysis identified several enriched motifs at predicted chromosome ends, including low-complexity AT-rich motifs and multiple motifs corresponding to rotations or reverse complements of the canonical insect telomeric repeat TTAGG. When equivalent TTAGG-like motifs were summarized together, clear terminal enrichment was detected at five chromosome ends: chr1-left, chr4-right, chrX2-left, chr5-right, and chr6-right (Supplementary Fig. S3). The strongest TTAGG-like signal occurred at chrX2-left, with 351 repeat counts per 10 kb window, followed by chr1-left, chr4-right, chr5-right, and chr6-right, with 226, 197, 111, and 68 repeat counts per 10 kb window, respectively. The remaining 15 chromosome ends showed weaker or no terminal enrichment for TTAGG-like motifs. This partial recovery of TTAGG-like terminal arrays is consistent with recent analyses of hemipteran chromosome assemblies, which support TTAGG as the predominant telomeric motif in Sternorrhyncha while also showing that telomeric repeats may be incompletely recovered at some assembled chromosome termini (Stoianova et al. 2025). The low-complexity motif AAAAT was also enriched near several chromosome ends, but because this motif is highly AT-rich and not clearly related to the canonical insect telomeric repeat, we interpret it conservatively as terminal or subtelomeric repetitive DNA rather than as evidence for an alternative telomere motif. Chromosome ends lacking strong TTAGG-like enrichment may reflect incomplete assembly through terminal repeat arrays, lower repeat copy number, or sequence divergence below the detection threshold used here. Terminal repeat motifs therefore provide independent support for the structure of several chromosome-scale scaffolds, but do not confirm telomere-to-telomere assembly for all predicted chromosomes.

### The *Pineus strobi* genome supports conservation of macrosynteny across sequenced adelgids

The chromosome-scale *P. strobi* assembly allowed us to test whether the conserved chromosomal linkage observed among *Adelges* genomes extends to *Pineus*, the *Pinus*-associated lineage recovered as sister to the remaining sampled adelgid lineages (Havill et al. 2007; Dial et al. 2023; Glendening et al. 2026). Across the ten chromosome-scale scaffolds of *P. strobi*, syntenic blocks largely connected genes to corresponding chromosome-scale scaffolds in *A. tsugae* and *A. abietis* (Fig. 2). Syntenic links involving the ten longest scaffolds of *A. cooleyi* were also broadly consistent with these major linkage groups, but the non-chromosome-scale status of the published *A. cooleyi* assembly limits inference about its precise chromosome structure. Although local gene order was disrupted by inversions and small-scale rearrangements, we observed little evidence for extensive interchromosomal reshuffling among major linkage groups. These results indicate that broad chromosomal linkage is conserved across the deepest sampled split within Adelgidae, extending patterns previously observed in *Adelges* to *Pineus*. However, chromosome-scale assemblies remain unavailable for several adelgid subgenera, particularly the monotypic subgenus *Pineodes*, represented by *P. pinifoliae*. The weak and variable placement of this species in existing limited-locus analyses makes it especially important for determining whether this pattern extends across the deepest splits in the family (Havill et al. 2025).

**Figure 2.**
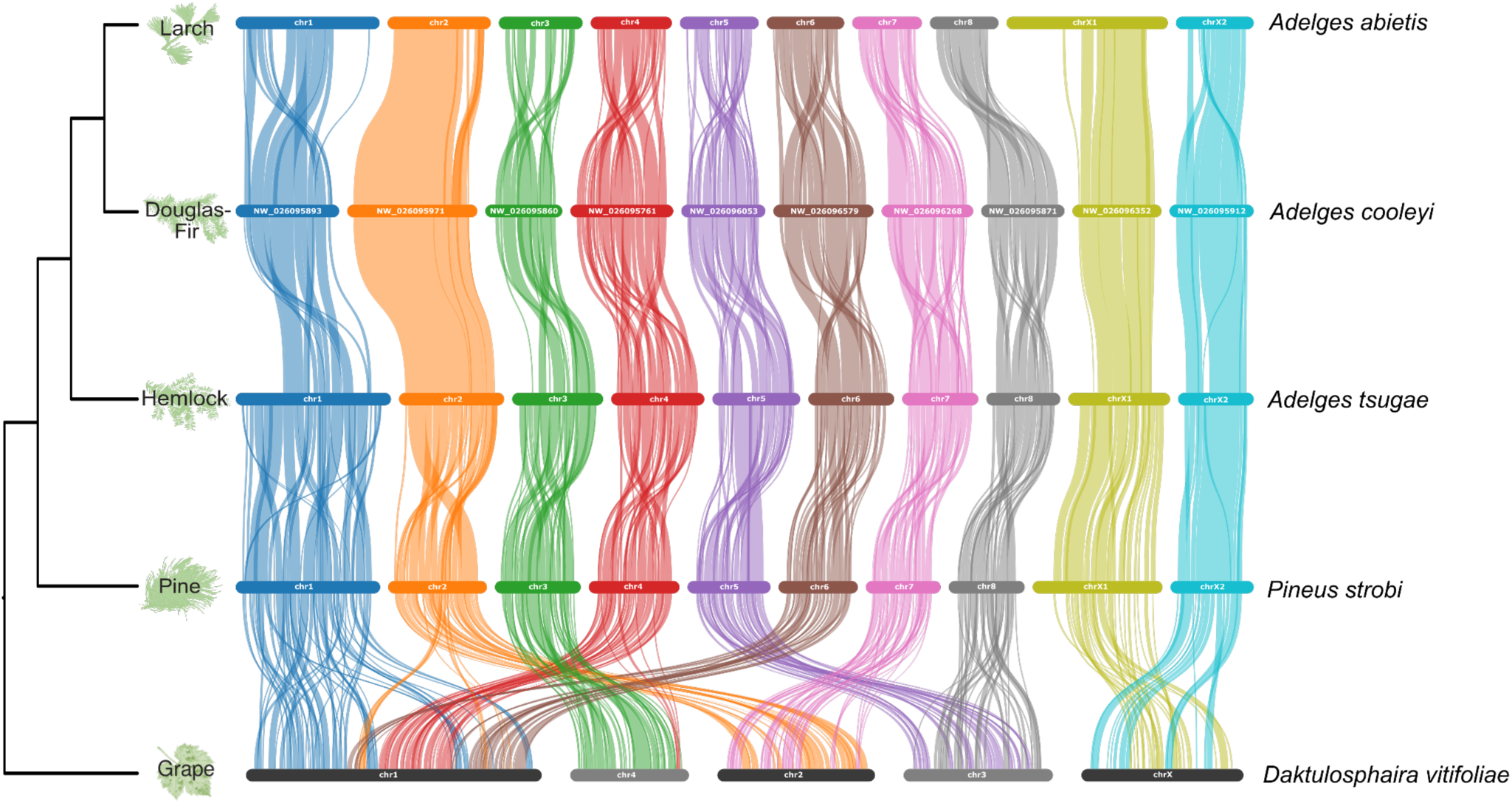
Conserved macrosynteny across adelgid and phylloxeran genomes. Comparative synteny plot showing the ten predicted chromosomal *Pineus strobi* scaffolds aligned with published genome assemblies of *Adelges abietis, A. cooleyi, A. tsugae*, and the grape phylloxera, *Daktulosphaira vitifoliae*. Host plants are indicated at branch tips. For *A. cooleyi*, which is not currently scaffolded to chromosome level, the ten longest scaffolds were selected for visualization. Horizontal bars represent chromosomes or major scaffolds, and colored ribbons connect collinear orthologous blocks between genomes. Each color corresponds to a major adelgid linkage group. The largely one-to-one correspondence of colored blocks among *P. strobi, A. tsugae, A. cooleyi,* and *A. abietis* indicates broad conservation of adelgid macrosynteny, whereas more dispersed connections to *D. vitifoliae* reflect the deep divergence between adelgids and phylloxerids.

Comparison with grape phylloxera, *Daktulosphaira vitifoliae*, extends this pattern to Phylloxeridae, the sister family to Adelgidae (Dial et al. 2023). The chromosome-scale assemblies of *P. strobi*, *A. tsugae*, and *A. abietis* each recover eight autosomes and two X chromosomes, whereas *D. vitifoliae* has four autosomes and a single X chromosome (Li et al. 2023). Despite this difference, each *D. vitifoliae* chromosome contains long syntenic blocks corresponding to a limited number of *P. strobi* chromosomes (Fig. 2). *Daktulosphaira vitifoliae* chr1 contains blocks corresponding primarily to *P. strobi* chr1, chr4, and chr6, together with a smaller portion of chr2. *Daktulosphaira vitifoliae* chr4 contains a largely collinear block corresponding to essentially all of *P. strobi* chr3, together with a smaller portion of chr4. *Daktulosphaira vitifoliae* chr2 corresponds primarily to *P. strobi* chr2 and chr7, whereas *D. vitifoliae* chr3 corresponds primarily to *P. strobi* chr5 and chr8. The single *D. vitifoliae* chrX likewise contains blocks corresponding to both *P. strobi* sex chromosomes, chrX1 and chrX2. Most *P. strobi* linkage groups therefore remain largely intact within a single *D. vitifoliae* chromosome, although *P. strobi* chr2 and chr4 each contribute blocks to two *D. vitifoliae* chromosomes. Although these comparisons cannot determine whether the difference in chromosome number arose through fusions in the phylloxerid lineage, fissions in the adelgid lineage, or a combination of both, the preservation of largely intact linkage groups indicates that the evolution of chromosome number was driven by a limited number of such events, with comparatively little exchange among major linkage groups, rather than the extensive interautosomal reshuffling observed in aphids (Li et al. 2023).

This conservation of major linkage groups in adelgids and grape phylloxera contrasts with patterns observed in aphids. Comparisons of chromosome-scale aphid genomes have revealed extensive interautosomal reshuffling in several lineages, particularly Macrosiphini, whereas X chromosome gene content has remained comparatively conserved (Mathers et al. 2021; Li et al. 2023; Huang et al. 2025). Holocentric chromosomes and cyclical parthenogenesis have both been proposed to facilitate the persistence of chromosome rearrangements because chromosome fragments can be stably inherited and rearranged karyotypes can propagate through clonal generations. However, these features alone are unlikely to explain the contrast. Holocentric chromosomes are widespread across Hemiptera, and cyclical parthenogenesis is ancestral to Aphidomorpha, yet available chromosome-scale comparisons show much stronger conservation of major linkage groups in adelgids and phylloxerids than in the most highly rearranged aphid lineages.

Phylogenetic depth is also unlikely to account for this contrast. Gene-order synteny generally decays with evolutionary distance in insects, but decay rates vary substantially among lineages, and major linkage groups can remain conserved over deep evolutionary timescales despite extensive local rearrangement (Van Dam et al. 2021; Wright et al. 2024). Although exact divergence times within Aphidomorpha remain uncertain, available adelgid and phylloxerid comparisons span evolutionary depths comparable to or greater than those within highly rearranged aphid clades. The split between *P. strobi* and the remaining sampled adelgid lineages has been estimated at approximately 90 Ma, older than the approximately 58 Ma crown age of the Aphidinae sampled by Huang et al. (2025) and substantially older than the divergences among the sampled Macrosiphini species, despite extensive autosomal reshuffling within that tribe (Havill et al. 2007; Huang et al. 2025). The split between Adelgidae and Phylloxeridae is deeper still. The greater conservation observed in the available adelgid and phylloxerid comparisons is therefore unlikely to be explained simply by less time for synteny to decay. Thus, the *P. strobi* genome supports the view that the extensive interautosomal reshuffling observed in several aphid lineages is not characteristic of Aphidomorpha as a whole but instead reflects lineage-specific genome dynamics within aphids.

Repeat dynamics provide one possible, nonexclusive explanation for this contrast. In aphids, transposable elements are enriched in autosomal synteny breakpoint regions, with LTR retrotransposons showing particularly strong enrichment, suggesting that repeat accumulation may have contributed to chromosomal rearrangement or marked regions prone to structural instability (Mathers et al. 2021). Aphid X chromosomes also show unusual repeat dynamics: although they exhibit long-term conservation of X-linked gene content, the X chromosomes of *M. persicae* and *A. pisum* are more repetitive than autosomes, and lineage-specific TE expansion has contributed substantially to X chromosome size differences (Mathers et al. 2021). In contrast, repetitive DNA accounts for a smaller fraction of the *P. strobi* genome, and LTR elements are rare (Table 1). Together with published adelgid genomes, this suggests that differences in repeat abundance, repeat distribution, or repeat activity may contribute to the different patterns of chromosome evolution observed between aphids and adelgids. Determining whether lower repeat loads or different repeat landscapes are consistently associated with conserved linkage groups will depend on more densely sampled chromosome-scale genomes across Adelgidae.

While chromosome-scale assemblies of *P. strobi*, *A. tsugae*, and *A. abietis* each recover ten major chromosomal scaffolds, and *A. cooleyi* is predicted to have 11 chromosomes (Steffan 1968), additional karyotype variation has been reported within Adelgidae (Gavrilov-Zimin et al. 2015). This pattern makes adelgids a useful system for studying chromosome evolution because most major linkage groups appear conserved, while some lineages still show differences in chromosome number or structure. As additional chromosome-scale genomes become available across pine, fir, larch, Douglas-fir, and hemlock adelgid lineages, it will become possible to identify lineage specific chromosome fusions, fissions, or sex chromosome changes and test whether these rearrangements are associated with repeat content, host specialization, or life-cycle transitions. Broader chromosome-scale sampling of phylloxerids and early-diverging aphid lineages will also allow rearrangement patterns to be compared across equivalent evolutionary timescales.

### The mitochondrial genome of *P. strobi* has large repeat and control regions and retains the ancestral insect gene order

We recovered a complete circular mitochondrial genome for *P. strobi* measuring 25,253 bp. The genome contains the expected insect mitochondrial gene complement, including 13 protein-coding genes, two rRNAs, and 22 tRNAs, and retains the ancestral insect mitochondrial gene order (Fig. 3). The *P. strobi* mitogenome is substantially larger than the 15–18 kb mitogenomes typical of insects and the approximately 15–20 kb mitogenomes reported for most aphids (Cameron 2014; Zhang et al. 2024), although a recent PacBio HiFi assembly recovered a 22.43 kb mitogenome for Tuberolachnus salignus (Crowley et al. 2024). As in the *A. cooleyi* and *A. tsugae* mitogenomes assembled with long-read data, much of the genome length is attributable to expanded noncoding regions (Dial et al. 2023; Brunet et al. 2025). In *P. strobi*, the repeat region spans 7.8 kb and the putative control region spans 2.7 kb. Comparisons with *A. tsugae* and *A. cooleyi* show broadly similar coding, rRNA, and tRNA content, but variation in the relative sizes of the repeat and control regions (Fig. 3). *Adelges cooleyi* has a 5.14 kb repeat region and a 4.31 kb control region (Dial et al. 2023), and *A. tsugae* has a 5.76 kb repeat region and a 4.66 kb control region (Brunet et al. 2025).

**Figure 3.**
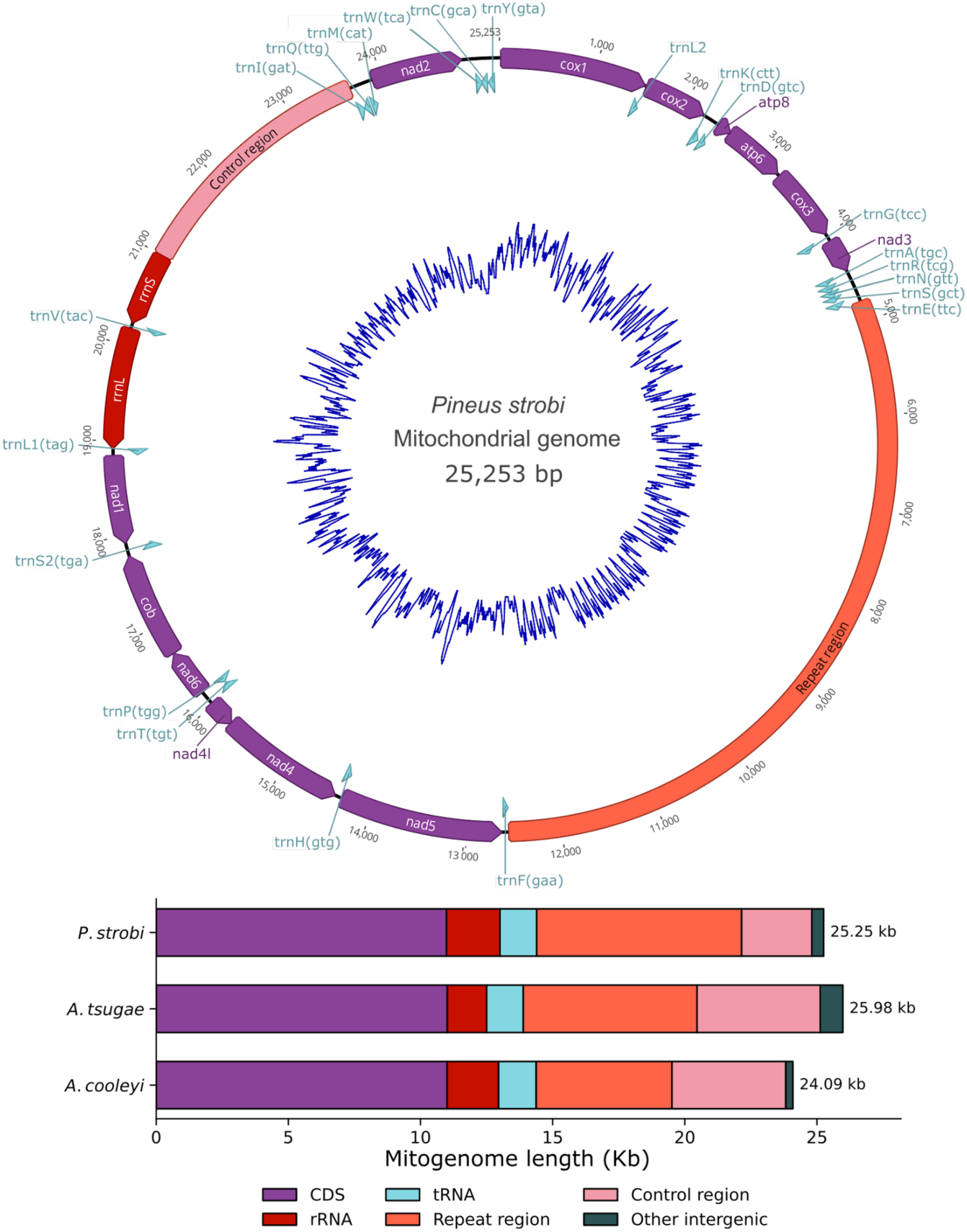
Mitochondrial genome organization of *Pineus strobi* and comparison with other adelgid mitogenomes. Circular map of the complete *P. strobi* mitochondrial genome, showing protein-coding genes, rRNAs, tRNAs, and major noncoding regions. Protein-coding genes are shown in purple, rRNAs in red, tRNAs in blue, the repeat region in orange, and the control region in pink. The inner blue trace shows GC content across the mitochondrial genome. The lower panel compares the proportional contribution of coding, RNA, repeat, control, and other intergenic sequence to mitogenome length in adelgids for which long read data is available, illustrating that large noncoding regions are a shared feature of available adelgid mitochondrial genomes.

Tandem Repeat Finder identified distinct tandem repeat arrays within both major noncoding regions of the *P. strobi* mitogenome. The repeat region is dominated by a 207 bp tandem repeat array spanning most of the region, whereas the putative control region contains a 283 bp tandem repeat array. BLAST searches of the 207 bp repeat unit recovered significant similarity primarily to aphidomorph mitochondrial sequences. Together, these observations are consistent with the interpretation that the aphidomorph mitochondrial repeat region predates the divergence of adelgids and aphids, while repeat expansion within the control region may be lineage specific.

The functional significance of the adelgid repeat region remains unresolved. Previous work proposed that the aphidomorph repeat region may have originated through ancient duplication of the control region and could represent a second replication-associated noncoding region (Song et al. 2019; Dial et al. 2023). The recovery of large repeat-rich regions in both *Pineus* and *Adelges* supports the view that expanded mitochondrial noncoding sequence is not restricted to a single adelgid lineage. Broader sampling of long-read aphidomorph mitogenomes would allow testing to determine the evolutionary origin, conservation, and functional relevance of these repeat-rich regions.

### The *Wolbachia* genome from *P. strobi* is within Supergroup A

In addition to the two bacteriocyte-associated gammaproteobacterial symbionts previously detected in *P. strobi* (Toenshoff et al. 2014), we recovered a complete circular genome of *Wolbachia* from the assembly. Mean HiFi read depth across the Wolbachia genome was 272x, comparable to mean depths of 283x and 328x across the two obligate symbiont genomes. To our knowledge, this represents the first complete *Wolbachia* genome recovered from an adelgid and the first evidence of *Wolbachia* infection in the family. Phylogenomic analysis of 94 *Wolbachia* genomes spanning the major described supergroups placed the *P. strobi* strain within Supergroup A, where it was most closely related to a *Wolbachia* strain from the thick-headed fly *Myopa testacea* (Fig. 4A). Whole-genome synteny comparisons between the *P. strobi* strain, its closest available relative from *M. testacea* isolate 54616 (Genbank Accession OZ034729.1), and a second *M. testacea*-associated strain, isolate 54618 (Genbank Accession OZ034748.1) revealed conserved local synteny interrupted by extensive rearrangement (Fig. 4B). This pattern is consistent with the dynamic genome architecture of *Wolbachia*, whose genomes commonly experience rearrangements associated with repetitive mobile elements, prophage WO regions, and homologous recombination (Vancaester and Blaxter 2023). The genome is 1,322,077 bp, encodes 1,396 predicted protein-coding genes, and has a GC content of 35.0% (Fig. 4C).

**Figure 4.**
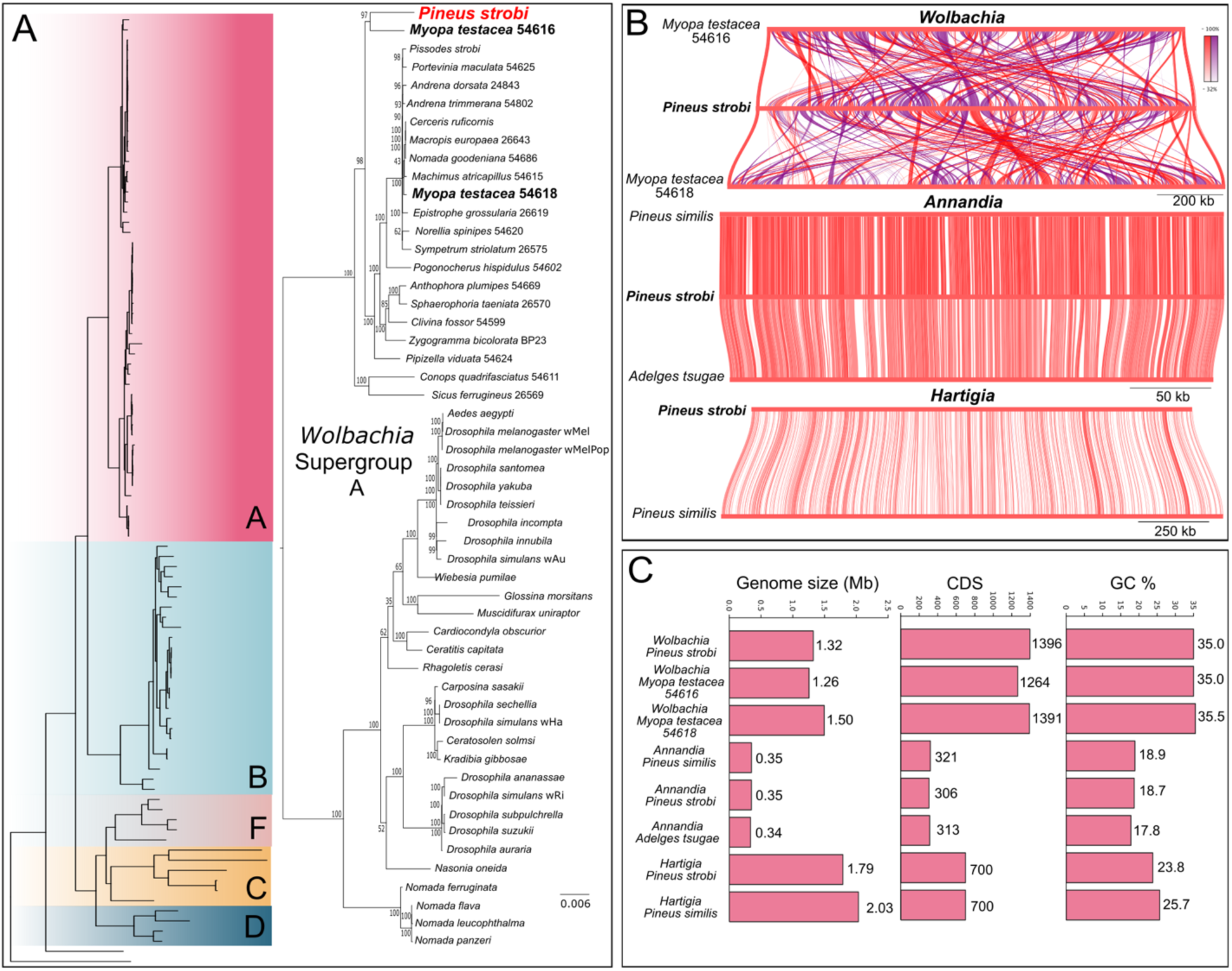
Phylogenetic placement and genome comparisons of bacterial symbionts associated with *Pineus strobi*. A) Phylogenetic placement of the *Wolbachia* genome from *P. strobi* within *Wolbachia* diversity. The broad tree at left derived from Vancaester and Blaxter 2023 and Palanichamy et al. 2025 shows representative *Wolbachia* supergroups, with the expanded Supergroup A tree shown at right, with taxa named for host species infected with *Wolbachia*. The *Wolbachia* strain from *P. strobi* falls within Supergroup A and is closely related to strains recovered from the fly *Myopa testacea* (Conopidae). B) Whole-genome alignment plots comparing the *P. strobi Wolbachia*, *Annandia*, and *Hartigia* genomes to their closest available relatives. Colored links connect homologous genomic regions, with darker colors indicating higher sequence similarity. Red links indicate genes in the same orientation, while purple links indicate inversions. C) Comparison of genome size, protein-coding gene number, and GC content among *P. strobi* symbionts and related bacterial symbiont genomes.

The recovery of a *Wolbachia* genome from *P. strobi* is notable in light of this species’ unusual reproductive biology relative to other adelgids. Typically, adelgid sexual reproduction is tightly coupled to host alternation: holocyclic lineages migrate to spruce, where the sexual generation and gall formation occur, whereas anholocyclic lineages remain on a single host and reproduce parthenogenetically (Havill and Foottit 2007). Some populations of *P. strobi* appear to retain a vestige of this ancestral host-alternating life cycle, as winged sexuparae can be produced and migrate to black spruce, but their offspring fail to complete development (Raske and Hudson 1964). Because *Wolbachia* is a maternally inherited endosymbiont known to alter arthropod reproduction through cytoplasmic incompatibility, male killing, feminization, and parthenogenesis induction (Werren et al. 2008), its presence in *P. strobi* raises the possibility that it is associated with the maintenance or reinforcement of parthenogenesis. However, the direction of causality remains unclear: *Wolbachia* could have contributed to disruption of the sexual phase, or it may have secondarily spread within an already anholocyclic lineage. Comparative screening of *Wolbachia* infection across *P. strobi* populations, related holocyclic *Pineus* species, and naturally or experimentally uninfected individuals will be needed to determine whether *Wolbachia* is functionally linked to the loss of a viable sexual generation. Future work could also closely examine genes related to host manipulation in adelgid and related *Wolbachia* strains.

### The obligate symbionts of *Pineus strobi* share perfect synteny with those from *Pineus similis*

Adelgids rely on paired obligate symbionts to supplement their nutritionally imbalanced sap diets. Genomic reconstructions indicate that, with probable contributions from the host, each symbiont consortium retains the capacity to complete biosynthetic pathways for all essential amino acids, although the division of these functions differs among adelgid lineages (Dial et al. 2022). We recovered complete circular genomes for the two bacteriocyte-associated nutritional symbionts previously identified from *P. strobi*: “*Candidatus Annandia pinicola*” (hereafter, *Annandia*) and “*Candidatus Hartigia pinicola*” (hereafter, *Hartigia*). Earlier work used microscopy, 16S/23S rRNA phylogenetics, and fluorescence in situ hybridization to show that *P. strobi* contains these two vertically transmitted gammaproteobacterial symbionts in distinct bacteriocyte types (Toenshoff et al. 2014), but genome sequences were not previously available for either symbiont. Prior to this study, the only sequenced obligate symbiont genomes from the *Pineus* genus were those of *Annandia* and *Hartigia* from *P. similis* (Dial et al. 2022). The *P. strobi Annandia* genome is highly reduced and similar in size and coding capacity to the *P. similis Annandia* genome and “*Candidatus Annandia adelgestsuga*” from *Adelges tsugae* (Weglarz et al. 2018), consistent with the interpretation that *Annandia* represents an ancient adelgid symbiont lineage retained across the deep split between *Pineus* and *Adelges* (Fig. 4B,C). Both *P. strobi* symbiont genomes retain perfect genome-wide synteny with their respective *P. similis* counterparts. Although the *Hartigia* genomes differ more substantially in total size, they contain very similar numbers of predicted protein coding genes (Fig. 4B,C), suggesting that their size difference largely reflects differential loss of noncoding intergenic DNA rather than major changes in gene content. Our metabolic reconstruction recovered the same predicted division of essential amino acid biosynthesis described for the *P. similis* pair: the partners are largely redundant for lysine biosynthesis, but *Annandia* encodes most other essential amino acid biosynthesis genes, including those for threonine, tryptophan, isoleucine, valine, arginine and leucine; *Hartigia* supplies the remaining step in phenylalanine biosynthesis and retains the histidine pathway and metE, encoding methionine synthase (Dial et al. 2022). The recovery of a complete *Wolbachia* genome from *P. strobi* therefore does not appear to reflect an additional obligate nutritional partner; instead, the conserved metabolic complementarity of *Annandia* and *Hartigia* supports their continued role as the primary nutritional symbionts of *P. strobi*.

## Conclusion

Together, the *Pineus strobi* nuclear, mitochondrial, and symbiont genomes expand the phylogenetic breadth of genomic resources available for Adelgidae and provide the first chromosome-scale reference for the genus *Pineus*. The chromosome-scale assembly supports broad conservation of major linkage groups across the deepest sampled split within Adelgidae. The recovery of a complete mitochondrial genome with expanded repeat-rich noncoding regions, complete genomes for the obligate nutritional symbionts *Annandia* and *Hartigia*, and the first reported *Wolbachia* genome from an adelgid further establishes *P. strobi* as a useful reference for studying genome architecture, phylogeny, mitochondrial genome evolution, and host-associated microbial partnerships. As additional chromosome-scale adelgid genomes become available, these resources will enable more detailed tests of how host specialization, life-cycle evolution, repeat dynamics, chromosome evolution, and symbiosis have shaped the diversification of Adelgidae.

## Materials and Methods

### Sample collection and species identification

Specimens of *P. strobi* were collected from bark on eastern white pine, *Pinus strobus,* at the Center for Invasive Insect and Disease Research of the USDA Forest Service in Hamden, Connecticut, USA, on 24 May 2020. These specimens were preserved in ethanol and used for PacBio HiFi sequencing. Species identity was determined based on host association, morphological characters, and mitochondrial cytochrome oxidase I (COI) barcoding. Voucher material was deposited at The Yale Peabody Museum under accession YPM#ENT996367. Additional live *P. strobi* specimens were collected from *Pinus strobus* on private property near Blue Ridge, Georgia, USA, on 30 June 2024 for Hi-C library preparation. These specimens were maintained alive on host material until processing. Because the Hi-C library was generated from an independently collected conspecific sample, we evaluated its compatibility with the PacBio HiFi assembly by assessing the alignment concordance of read mapping and Hi-C contact patterns (see Results and Discussion). Separate *P. strobi* specimens that had been maintained at Cornell University were used for flow cytometric estimation of genome size at Texas A&M University in 2019.

### DNA extraction, library prep, and sequencing

PacBio HiFi and Hi-C libraries were prepared in-house. High-molecular-weight DNA was extracted from pooled whole *P. strobi* individuals using the Quick-DNA HMW MagBead Kit (Zymo Research, Irvine, CA, USA; cat. no. D6060), with modifications for low-input material, including scaled down bead and elution volumes and extended incubation/mixing steps to improve DNA yield and fragment length. DNA quantity and quality were assessed using Qubit fluorometry and NanoDrop spectrophotometry (Thermo Fisher Scientific, Waltham, MA, USA), and fragment size distributions were assessed using an Agilent Fragment Analyzer system (Agilent Technologies, Santa Clara, CA, USA). A PacBio HiFi library was prepared using the SMRTbell Prep Kit 3.0 (Pacific Biosciences, Menlo Park, CA, USA) following the manufacturer’s protocol and sequenced on a PacBio Revio system using SPRQ chemistry. For Hi-C sequencing, live *P. strobi* individuals were flash frozen in liquid nitrogen and immediately processed using the Arima-HiC+ kit (Arima Genomics, Carlsbad, CA, USA) following the manufacturer’s user guide for animal tissues. The resulting Hi-C library was sequenced on an Illumina NovaSeq X platform using 150 bp paired-end sequencing. All sequencing was performed at the Yale Center for Genome Analysis (New Haven, CT).

### Genome assembly, contaminant filtering, and scaffolding

BAM files of PacBio HiFi reads were converted to FASTQ format with samtools v1.21 and assembled with hifiasm v0.25.0 (Li et al. 2009; Cheng et al. 2021). The initial assembly was screened for non-target sequences using BlobTools v1 based on read coverage, GC content, and taxonomic assignment from DIAMOND searches against the NCBI nonredundant protein database (Laetsch and Blaxter 2017; Buchfink et al. 2021). Contigs assigned to non-insect categories were removed, except for 77 contigs totaling 3.91 Mb labeled as “no-hit” by BlobTools that were retained because they lacked taxonomic evidence for contaminant origin. HiFi reads were also assembled with metaFlye v2.9.5 to recover circular symbiont and mitochondrial genomes (Kolmogorov et al. 2020). Hi-C reads were trimmed with Trimmomatic v0.39 using default settings (Bolger et al. 2014) and mapped to the decontaminated nuclear contigs with the Arima mapping pipeline (https://github.com/arimagenomics/mapping_pipeline). Briefly, reads were aligned to the assembly with BWA-MEM (Li and Durbin 2009), filtered to retain informative 5′ alignment positions, paired, filtered by mapping quality, sorted, and deduplicated before downstream Hi-C scaffolding. Because the Hi-C library was generated from an independently collected conspecific sample, we evaluated compatibility between the Hi-C reads and the PacBio HiFi assembly using mapping statistics, alignment concordance, and Hi-C contact patterns. The Hi-C alignments were then used for chromosome-scale scaffolding with YaHS v1.2.2 (Zhou et al. 2023). Scaffolding output was inspected with Juicebox, and the final assembly was manually reviewed using the Hi-C contact map (Durand et al. 2016). Genome size and heterozygosity were estimated from 31-mer frequencies in the PacBio HiFi reads using meryl v1.4.1 and GenomeScope2 (Ranallo-Benavidez et al. 2020; Rhie et al. 2020). An independent estimate of genome size was obtained by flow cytometry following standard protocols for insects (Johnston et al. 2019). Consensus quality and k-mer completeness of the cleaned assembly were evaluated with Merqury v.1.3 (Rhie et al. 2020) using the same 31-mer database derived from the HiFi reads. Assembly completeness was assessed with BUSCO v6.0.0 using the Hemiptera odb10 dataset (Tegenfeldt et al. 2025).

### Gene prediction

Gene annotation of the *P. strobi* nuclear genome was performed with EGAPx, the public implementation of the NCBI Eukaryotic Genome Annotation Pipeline (https://github.com/ncbi/egapx). EGAPx performs repeat masking internally with WindowMasker before transcript and protein evidence alignment (Morgulis et al. 2006). Publicly available paired-end Illumina RNA-seq reads from *Pineus* sp. PispAD (SRR5134714) were used as transcript evidence. In a previous phylogenomic analysis, we recovered a full-length COI transcript from this dataset that matched *P. strobi* by 99.85%, supporting its use as transcriptomic evidence for annotation (Dial et al. 2023). Using the supplied *P. strobi* NCBI Taxonomy ID, EGAPx selected protein and HMM evidence sets from related insects for gene prediction, including *Myzus persicae*, *Aphis gossypii*, *Rhopalosiphum padi*, *Lycorma delicatula*, *Planococcus citri*, *Rhodnius prolixus*, *Bemisia tabaci*, *Cimex lectularius*, *Drosophila melanogaster*, and *Homalodisca vitripennis*. RNA-seq reads were aligned with STAR (Dobin et al. 2013), protein evidence was aligned using miniprot (Li 2023), and transcript and protein evidence were integrated by Gnomon (Souvorov et al. 2010) to predict gene models. Prediction of noncoding RNA features was performed with tRNAscan-SE (Chan et al. 2021) and cmsearch/Infernal searches against Rfam (Nawrocki and Eddy 2013; Ontiveros-Palacios et al. 2025). Functional annotations were assigned by EGAPx using model quality, homology, and orthology information. Proteome completeness was assessed within EGAPx using BUSCO v5.7.1 with the Hemiptera odb10 dataset (Manni et al. 2021) with the longest isoform per gene. The mitochondrial genome was annotated with mitoZ v3.6 (Meng et al. 2019), and endosymbiont genomes were annotated with Prokka v1.14.6 (Seemann 2014).

### Repeat content and telomere annotation

Repetitive elements were characterized independently of the EGAPx gene annotation workflow. A de novo repeat library was generated with RepeatModeler2 v2.0.5 (Flynn et al. 2020) and used to annotate repetitive elements in the *P. strobi* nuclear genome with RepeatMasker v3.6 (Tarailo-Graovac and Chen 2009). Repeat annotations were summarized by repeat class to estimate the genomic contribution of DNA transposons, LINEs, LTRs, SINEs, simple repeats, low-complexity sequence, and unclassified repeat families. Because EGAPx uses WindowMasker internally for gene prediction rather than repeat-family classification, RepeatModeler2 and RepeatMasker annotations were used for all reported repeat content estimates. Tandem Repeat Finder (Benson 1999) was used to identify and quantify repeats in the control and repeat regions of the mitochondrial genome.

Candidate telomeric and terminal repeat motifs were identified with tidk v0.2.65 (Brown et al. 2025). We first used the tidk explore function to identify enriched terminal repeat motifs using repeat units of 4–15 bp, a terminal search distance of 0.01, and a minimum threshold of 30 repeat copies. This search recovered a TTAGG-like motif, corresponding to the ancestral insect telomeric repeat, as well as additional terminally enriched motifs, including AAAAT and AGCGCC. Candidate motifs were then quantified genome-wide with tidk search in 10 kb windows, with forward and reverse counts summed for each window. Because tidk reports tandem repeats in canonicalized form, rotations and reverse complements of the ancestral insect telomeric repeat TTAGG were treated as equivalent. For each 10 kb window, the TTAGG-like repeat signal was defined as the highest count observed among these equivalent motifs. For each chromosome-scale pseudomolecule, terminal repeat abundance was calculated as the maximum repeat count detected in any 10 kb window within the last 100 kb of each chromosome end. Chromosome ends with more than 40 TTAGG-like repeats per 10 kb window were considered strongly enriched. Additional AAAAT and AGCGCC signals were plotted as terminally enriched repeat candidates separately from TTAGG-like telomeric enrichment.

### Synteny analysis

To assess broad patterns of macrosynteny across adelgids, we compared predicted protein sequences from *P. strobi* with available high-quality adelgid and phylloxera genome assemblies. For each species, the longest isoform per gene was retained for analysis. Pairwise protein similarity searches from all vs. all DIAMOND BLASTp searches were used as input for MCScanX v1.0.0 (Wang et al. 2012) to identify collinear gene blocks. For *A. cooleyi*, which does not yet have a published chromosome-scale assembly, visualization was restricted to the ten longest scaffolds representing 63.2% of the total assembly length to provide a cautious comparison of large-scale linkage patterns rather than a definitive chromosome-level reconstruction. Synteny plots were generated with SynVisio (Bandi and Gutwin 2020) and edited in Inkscape for clarity. Synteny plots for endosymbiont and mitochondrial genomes were generated with pyGenomeViz v1.6.1 (Shimoyama 2024) using MMseqs2 v18-8cc5c (Steinegger and Söding 2017) for sequence alignment.

### Wolbachia phylogenomic analysis

To determine the phylogenetic placement of the *P. strobi Wolbachia* genome, taxa were selected for representation across each major *Wolbachia* Supergroup, using recent studies on *Wolbachia* genomic diversity as references (Vancaester and Blaxter 2023; Palanichamy et al. 2025), and the top 20 hits in a blastn search of the 16S sequence from the *Wolbachia* of *P. strobi* against the NCBI nucleotide (nt) database were selected. Predicted protein sequences from the newly assembled *Wolbachia* genome and a total of 93 publicly available *Wolbachia* genomes were compared using OrthoFinder v3.1.3 (Emms and Kelly 2019). A *Wolbachia* phylogeny was inferred from the OrthoFinder concatenated species tree alignment, generated from near-single-copy orthogroups under the MSA workflow with mafft v7.505 (Katoh and Standley 2013) for alignment. The resulting 53,305 amino-acid alignment was analyzed with IQ-TREE v2.03 (Minh et al. 2020) under the Q.BIRD+F+I+R4 model with 1,000 ultrafast bootstrap replicates and 1,000 SH-aLRT replicates. The resulting tree was visualized with FigTree (Rambaut 2018) and edited in Inkscape.

## Data Availability

The raw PacBio HiFi and Hi-C sequencing reads, nuclear genome assembly and annotation, complete mitochondrial genome, and complete symbiont genome assemblies generated in this study will be deposited in NCBI. BioProject and individual accession numbers are pending and will be added during peer review and before publication.

## Acknowledgements

We thank the Yale Center for Genome Analysis for sequencing libraries and J. Spencer Johnston (Department of Entomology, Texas A&M University, College Station, Texas, USA) for performing flow-cytometric genome size estimation. Computational analyses were performed using the Deigo high-performance computing cluster at the Okinawa Institute of Science and Technology (OIST).

